# Circadian rhythms tied to changes in brain morphology in a densely-sampled male

**DOI:** 10.1101/2024.04.10.588906

**Authors:** Elle M. Murata, Laura Pritschet, Hannah Grotzinger, Caitlin M. Taylor, Emily G. Jacobs

## Abstract

Circadian, infradian, and seasonal changes in steroid hormone secretion have been tied to changes in brain volume in several mammalian species. However, the relationship between circadian changes in steroid hormone production and rhythmic changes in brain morphology in humans is largely unknown. Here, we examined the relationship between diurnal fluctuations in steroid hormones and multiscale brain morphology in a precision imaging study of a male who completed forty MRI and serological assessments at 7 A.M. and 8 P.M. over the course of a month, targeting hormone concentrations at their peak and nadir. Diurnal fluctuations in steroid hormones were tied to pronounced changes in global and regional brain morphology. From morning to evening, total brain volume, gray matter volume, and cortical thickness decreased, coincident with decreases in steroid hormone concentrations (testosterone, estradiol, and cortisol). In parallel, cerebrospinal fluid and ventricle size increased from A.M. to P.M. Global changes were driven by decreases within the occipital and parietal cortices. These findings highlight natural rhythms in brain morphology that keep time with the diurnal ebb and flow of steroid hormones.

**Significance Statement:** Though rhythmic changes in steroid hormone secretion have been tied to changes in brain volume in several mammalian species, this relationship has not been well characterized in humans. In this precision neuroimaging study, we found that global and regional brain morphology and steroid hormone levels exhibit tandem circadian rhythms. These findings provide high-resolution insight into the anatomical signature of diurnal changes in brain morphology and steroid hormone production in a human male and reveal the metronomic regularity of these rhythms over time.

## Introduction

Endogenous biological rhythms guide physiological processes throughout the body in nearly all organisms. Some of the most pronounced biological rhythms are the time-varying secretion of steroid hormones. Testosterone, estradiol, and cortisol exhibit circadian rhythms, with production peaking in the morning and decreasing throughout the day (Ankarberg-Lindgren & Norjavaara, 2008; Diver et al., 2003; Pruessner et al., 1997; Rose et al., 1972), yet the influence of this diurnal ebb and flow of hormone production on the human brain is remarkably understudied. Several lines of evidence suggest such a link exists. Rhythmic changes in steroid hormone secretion, often marked by increased levels in breeding season and reduced levels in winter (Prendergast et al., 2002), have been tied to changes in brain morphology in animal models. In bank voles, brain and hippocampal mass are elevated during the summer and reduced in winter months (Yaskin, 1984). Mice exhibit decreased total brain size and hippocampal volume (Pyter et al., 2005) and hamsters show reduced soma size in the hippocampus (Workman et al., 2011) in short versus long photoperiods. Songbirds also exhibit seasonal fluctuations in brain structure, morphological changes that coincide with changes in steroid hormone production (Bottjer et al., 1986; DeVoogd & Nottebohm, 1981; Nottebohm, 1981). Human MRI studies have begun to characterize the role of circadian rhythms in modulating brain structure and function. There are reports of diurnal differences in functional connectivity (Farahani et al., 2021; Orban et al., 2020), cerebral blood flow (Elvsåshagen et al., 2019), water diffusivity (Thomas et al., 2018; Tuura et al., 2021), and brain volume (Karch et al., 2019; Trefler et al., 2016; Zareba et al., 2023, 2024). However, the association between circadian changes in hormone production and rhythmic changes in the brain is unclear.

Precision neuroimaging studies have begun to advance the study of brain–hormone relationships by densely scanning individuals with high temporal resolution (Jacobs, 2023; Pritschet et al., 2021; Pritschet & Jacobs, 2023). For example, our group found that rhythmic changes in steroid hormones across the 24-hour diurnal cycle (Grotzinger et al., 2023) and 30-day menstrual cycle (Fitzgerald et al., 2020; Mueller et al., 2021; Pritschet et al., 2020) are sufficient to drive changes in the brain’s functional network architecture. Changes in brain morphology are also evident across these timescales. Subregions of the medial temporal lobe wax and wane in step with endocrine changes across the menstrual cycle (Zsido et al., 2023), an effect that is eliminated by chronic hormone suppression (Taylor et al., 2020). Precision imaging studies are yielding new insights into steroid hormones as rapid modulators of brain plasticity, in keeping with thirty years of evidence from animal models (Taxier et al., 2020). To date, we lack a comparable understanding of how rhythmic endocrine changes influence brain morphology across shorter cycles, such as the diurnal change in hormone production.

To determine the influence of diurnal hormone fluctuations on brain morphology, we used a dense-sampling design to collect MRI, serum, saliva, and mood data from a healthy adult male every 12–24 hours for 30 consecutive days. Time-locked sessions occurred at 7 A.M. and 8 P.M., capturing the peak and nadir of steroid hormone production. By metronomically varying the timing of MRI and biofluid collection, we were able to track the diurnal ebb and flow of endocrine and neural changes in the human brain.

Diurnal fluctuations in steroid hormones were tied to pronounced changes in global and regional brain morphology. From morning to evening, total measures of brain volume, gray matter volume (GMV), and cortical thickness (CT) decreased significantly, paralleling decreases in testosterone, estradiol, and cortisol. Cerebrospinal fluid (CSF) volume and ventricle size showed the opposite pattern, increasing throughout the day. At the regional level, cortical reductions were most apparent in visual cortices, including the striate and extrastriate cortex; subcortical reductions were evident in the cerebellum, brain stem, and right hippocampus. These findings demonstrate a natural rhythm to brain morphology, a cadence that pulses in time with the diurnal ebb and flow of steroid hormones.

## Materials and Methods

### Participant

The participant was a healthy 26-year-old right-handed Caucasian male with no history of neuropsychiatric diagnosis, endocrine disorders, or prior head trauma. The participant gave written informed consent for a study approved by the University of California, Santa Barbara Human Subjects Committee and was paid for participation in the study.

### Experimental design

The methods for this study are reported in Grotzinger et al., 2023. Briefly, the participant underwent venipuncture and brain imaging every 12–24 hours for 30 consecutive days. At each session the participant completed a questionnaire to assess stress, sleep, and mood states, followed by endocrine sampling at 7 A.M. (morning sessions) and/or at 8 P.M. (evening sessions). The participant gave a 2mL saliva sample at each session, followed by a blood sample. On days with two sessions, the participant underwent one blood draw per day (**Fig. 1A**), per safety guidelines. Morning endocrine samples were collected after at least 8 hours of overnight fasting, and evening endocrine samples were collected following 1.5 hours of abstaining from consumption of food or drink (excluding water). The participant refrained from consuming caffeinated beverages before each morning session.

**Figure 1.**
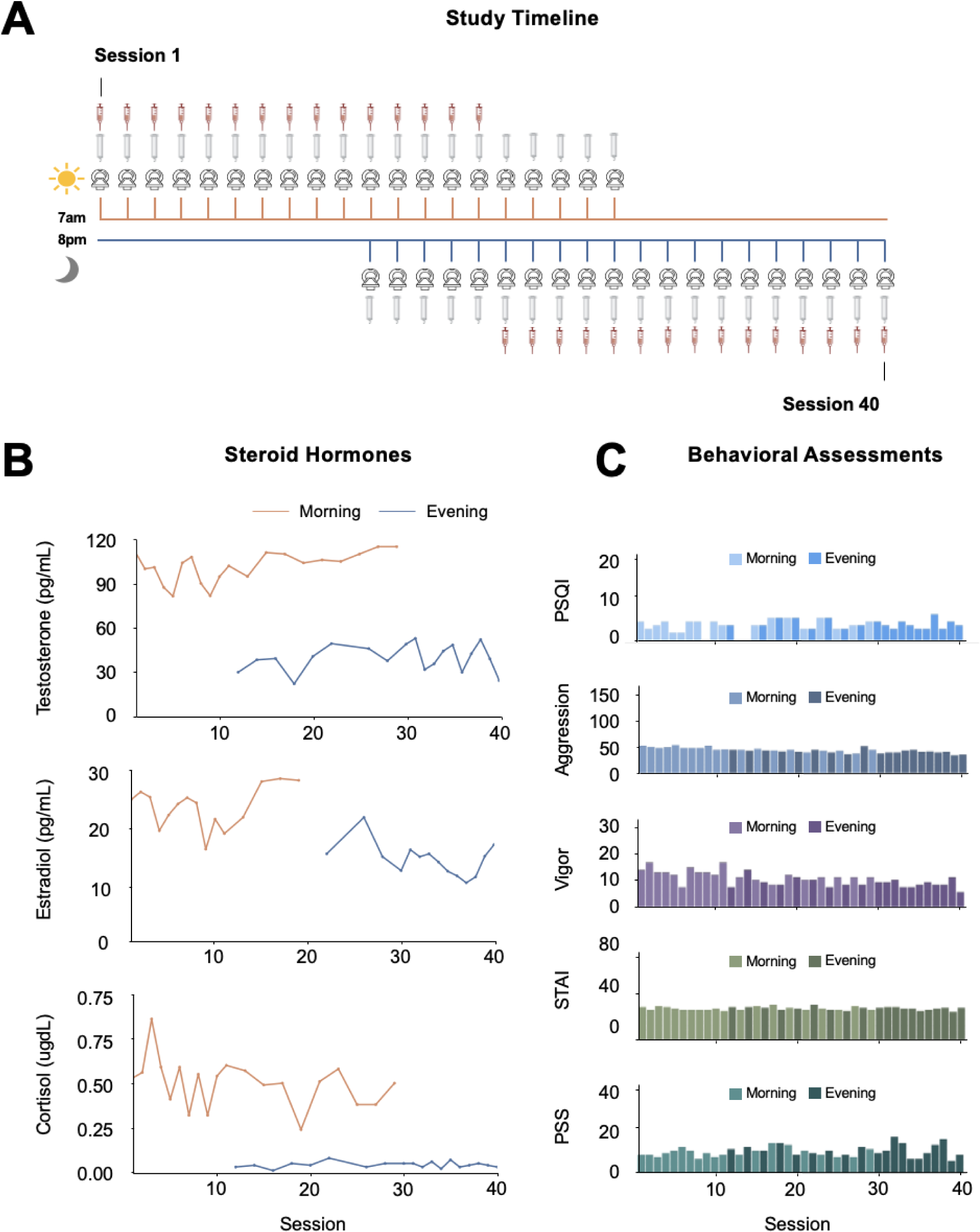
Precision imaging study design. (**A**) For 30 consecutive days, a healthy 26-year-old male underwent brain imaging and serological assessment every 12-24 hours for a total of 40 sessions. **(B)** Steroid hormones demonstrated typical diurnal patterns, with testosterone (saliva), estradiol (serum), and cortisol (saliva) peaking in the morning and significantly reduced in the evening. **(C)** The subject completed mood assessments (e.g., sleep quality, mood, anxiety, stress) at each session. All measures were within the standard range and remained largely consistent across the study duration. Abbreviations: PSQI= Pittsburgh Sleep Quality Index; STAI = State-Trait Anxiety Inventory; PSS = Perceived Stress Scale.

#### Behavioral assessments

The following scales (adapted to reflect the past 12–24 hours) were administered at each session as part of the questionnaire: the Perceived Stress Scale (PSS) (Cohen et al., 1983), Pittsburgh Sleep Quality Index (PSQI) (Buysse et al., 1989), State-Trait Anxiety Inventory for Adults (STAI) (Speilberger & Vagg, 1984), Profile of Mood States (POMS) (Pollock et al., 1979), and Aggression Questionnaire (Buss & Perry, 1992). The questionnaire for the evening sessions excluded the PSQI to avoid redundancy. All mood measures fell within standard reference ranges (**Figure 1C**).

#### Endocrine procedures

A ∼2 mL saliva sample was obtained via passive drooling into a wide-mouthed plastic cryovial. The participant refrained from eating and drinking (besides water) at least 1.5 hours before saliva sample collection, and the morning samples were collected after fasting overnight. The participant pooled saliva for 5–10 minutes before depositing the sample into the cryovial to determine total testosterone and cortisol concentrations. The sample was stored at –20°C until assayed. Saliva concentrations were determined via enzyme immunoassay at Brigham and Women’s Hospital Research Assay Core.

Immediately after the saliva sample was obtained, a licensed phlebotomist inserted a saline-lock intravenous line into either the dominant or non-dominant hand or forearm daily to evaluate total testosterone, free testosterone, cortisol, and 17β-estradiol concentrations. One 10 mL blood sample was collected in a vacutainer SST (BD Diagnostic Systems) each session. The sample clotted at room temperature for ∼60 min until centrifugation (2100 x g for 10 min) and was then aliquoted into three 2 mL microtubes. Serum samples were stored at –20°C until assayed. Serum concentrations were determined via liquid chromatography mass-spectrometry at the Brigham and Women’s Hospital Research Assay Core. Assay sensitivities, dynamic range, and intra-assay coefficients of variation (respectively) were as follows: estradiol, 1 pg/mL, 1–500 pg/mL, < 5% relative standard deviation (*RSD*); testosterone, 1.0 ng/dL, 1–200 ng/dL, <2% *RSD*; cortisol, 0.5 ng/mL, 0.5–250 pg/mL, <8% *RSD*. Note that estradiol and free testosterone measurements were acquired from serum samples, resulting in 30 timepoints.

### MRI acquisition

At each session, the participant underwent a whole-brain structural MRI and 15-minute eyes-open resting-state scan conducted on a Siemens 3T Prisma scanner equipped with a 64-channel phased-array head coil. High-resolution anatomical scans were acquired using a T1-weighted magnetization prepared rapid gradient echo (MPRAGE) sequence (TR = 2500 ms, TE = 2.31 ms, TI = 934 ms, flip angle = 7°, 0.8 mm thickness), followed by a gradient echo fieldmap (TR = 758 ms, TE1 = 4.92 ms, TE2 = 7.38 ms, flip angle = 60°). A T2-weighted turbo spin echo (TSE) scan was also acquired with an oblique coronal orientation positioned orthogonally to the main axis of the hippocampus (TR/TE = 8100/50 ms, flip angle = 122°, 0.4 × 0.4 mm^2^ in-plane resolution, 2 mm slice thickness, 31 interleaved slices with no gap, total acquisition time = 4:21 min). Resting-state functional MRI scans allowed for indirect assessments of head motion (Grotzinger et al., 2023).

### Whole-brain morphology assessments

Image processing for evaluating GMV, white matter volume (WMV), and CT was conducted with Advanced Normalization Tools (ANTs, version 2.1.0). A subject-specific anatomical template (SST) was created with multivariate template construction. Subject-specific tissue priors were created using the OASIS population template priors. Structural images from each session were then registered to this template and processed with the ANTs CT pipeline (Tustison et al., 2014). This pipeline consists of N4 bias correction, tissue segmentation, and cortical thickness estimation. N4 bias correction minimizes field inhomogeneity effects. Tissue segmentation via Atropos makes tissue segmentation estimates of white matter, gray matter, deep gray matter, and cerebrospinal fluid by using prior knowledge to guide segmentation. The Diffeomorphic Registration-based Cortical Thickness (DiReCT) algorithm was used to evaluate CT. CT measures were created based on diffeomorphic mappings between gray matter–white matter interface and the estimated gray matter–cerebrospinal fluid interface (Das et al., 2009). Gray matter measures were created by first normalizing each gray matter tissue mask to the template. Then, it was multiplied to a Jacobian image computed by affine and non-linear transforms. Brain volume and CT summary measures were output automatically for each scan session. The 400-region Schaefer parcellation was first warped to the SST then applied to volumetric images to assess regional level measures of CT and GMV by calculating the first eigenvariate across all voxels within each parcel (Schaefer et al., 2018). Regional-level metrics were computed by averaging parcels following the 42-region Schaefer scheme (Schaefer et al., 2018; Yeo et al., 2011). Measures of CSF, ventricle size, and subcortical volume were calculated by running the T1 images through the longitudinal recon-all pipeline from Freesurfer (Dale et al., 1999) and segmented into the 28 regions-of-interest determined via the aseg parcellation (Fischl et al., 2002).

### Medial temporal lobe subfield assessments

High resolution medial temporal lobe volumes (CA1, CA2/3, dentate gyrus, subiculum, perirhinal, entorhinal and parahippocampal cortex) were measured from a dedicated T2-weighted image of the hippocampi, co-registered to a T1-weighted MRI using the automatic segmentation of hippocampal subfields (ASHS) package (Yushkevich et al., 2015) and the Princeton Young Adult 3T ASHS Atlas (n = 24, mean age 22.5 years) (Aly & Turk-Browne, 2015). T2w scans and segmentations were first visually examined using ITK-SNAP (Yushkevich et al., 2006) for quality assurance and then subjected to manual editing in native space using ITK-SNAP (v.3.8.0-b; author CMT). Boundaries between perirhinal, entorhinal, and parahippocampal cortices were established in keeping with the Olsen-Amaral-Palombo (OAP) segmentation protocol (Palombo et al., 2013). In instances where automatic segmentation did not clearly correspond to the underlying neuroanatomy, such as when a certain label was missing several gray matter voxels, manual retouching allowed for individual voxels to be added or removed. All results are reported using the manually retouched subregion volumes to ensure the most faithful representation of the underlying neuroanatomy. Scans were randomized and segmentation was performed in a random order.

### Statistical analyses

All analyses were conducted in R (version 4.3.2). Independent samples t-tests were run to compare global and regional brain structure measures between morning and evening sessions. T-test results were considered statistically significant if they met the Bonferroni correction for multiple comparisons: *p* < .0056 (*p* < .05/9 measures) for global morphology measures, *p* < .007 (*p* < .05/7 regions) for medial temporal subfields, *p* < .001 (*p* < .05/42 regions) for regional cortical measures, and *p* <. 0.0018 (*p* <.05/28 regions) for regional subcortical measures. Effect size was estimated using Cohen’s *d*. To evaluate the impact of time of day (TOD) on GMV and CT in individual nodes, multivariate regressions were performed (false discovery rate [FDR]-corrected). Additionally, correlation matrices between brain measures and steroid hormones (testosterone [saliva], estradiol [serum], cortisol [saliva]) were computed and Bonferroni-corrected. Because most steroid hormones (testosterone, cortisol) did not display a normal distribution (Shapiro-Wilk test: *p* < .05), non-parametric Spearman’s rank correlations were computed. To note, session 24 was excluded from analyses due to an abnormal dip in testosterone after the participant received the COVID-19 vaccine. In this evening session, total testosterone (saliva) was 6.54 pg/mL, which was ∼83 % lower than the average evening testosterone level (M=37.93 pg/mL).

## Results

Evaluations of steroid hormones confirmed the expected peak of testosterone, estradiol, and cortisol in the morning and nadir in the evening. Testosterone, estradiol, and cortisol decreased from morning to evening by ∼61%, ∼38%, and ∼92%, respectively (**Figure 1B**, **Table 1-1**).

### Global brain morphology by time of day and steroid hormone levels

Changes in brain morphology were evident by TOD. Total brain volume (*t*(29.37) = –6.93, *p* < .001, *d* = –2.23) and total GMV (*t*(35.70) = –7.22, *p* < .001, *d* = –2.32) peaked in the morning and dipped in the evening. There was no observable difference in WMV at a corrected significance threshold (*t*(34.33) = –2.32, *p* = .026, *d* = –0.75) (**Figure 2A**, **Table 2-1**). In contrast, global increases were observed in CSF (*t*(34.00) = 3.53, *p* = .001, *d* = 1.14), lateral ventricles (*t*(33.27) = 3.72, *p* < .001, *d* = 1.97) and 4th ventricle (*t*(27.68) = 3.60, *p* = .001, *d* = 1.16) volumes from morning to evening (**Figure 2B**, **Table 2-1**). With the exception of WMV, Cohen’s *d* values indicated a large effect of TOD on global brain morphology measures. These relationships paralleled diurnal changes in steroid hormone concentrations. Steroid hormones demonstrated a positive correlation with total brain volume (testosterone: *r* = 0.58, *p* < .001, estradiol: *r* = 0.70, *p* < .001, cortisol: *r* = 0.56, *p* < .001) and global GMV (testosterone: *r* = 0.67, *p* < .001, estradiol: *r* = 0.69, *p* < .001, cortisol: *r* = 0.61, *p* < .001, Bonferroni corrected). Global cortical thickness demonstrated a comparable decrease from morning to evening, with the same positive relationship with all three hormones (**Table 2-1).** With the exception of aggression and total brain volume (*r* = 0.55, *p* < .001), there were no significant associations between global brain morphology measures and mood assessments (**Figure 2-1**).

**Figure 2.**
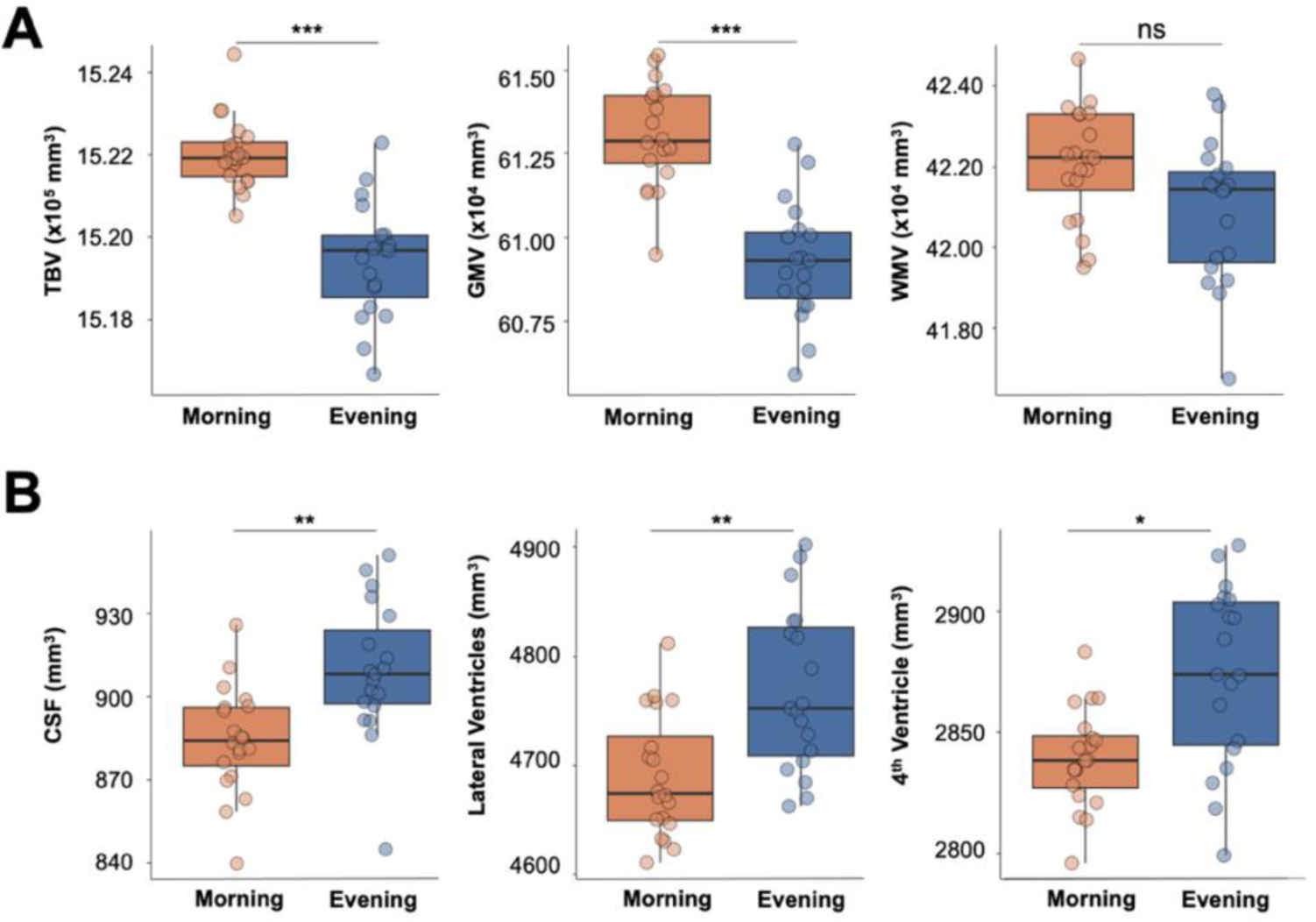
Neuroanatomical changes throughout the day in a densely-sampled male. (**A**) Total brain volume and gray matter decreased over the course of the day, while white matter volume did not (at a Bonferroni corrected threshold). **(B)** Cerebrospinal fluid, lateral ventricle, and fourth ventricle volume all increased from morning to evening. Abbreviations: TBV = Total Brain Volume; GMV = Gray Matter Volume; WMV = White Matter Volume; CSF = Cerebrospinal Fluid; ns = not significant, \**p* < .006, \*\**p* < .001, *** *p* <.0001.

### Regional gray matter volume is tied to time of day and steroid hormone concentrations

To investigate whether changes in morphology were ubiquitous throughout the brain, a regional approach was taken to pinpoint areas most sensitive to diurnal rhythms. Pronounced diurnal decreases in GMV were evident in the parietal and occipital cortices, including extrastriate and striate, temporal occipital, and precuneus regions. The effect size was large across regions (all *d* < –1.33, *p* < .001, **Figure 3**, **Table 3-1**). The magnitude of these regional reductions was strongly associated with the diurnal reduction in steroid hormones (**Table 1**, **Table 1-2**). Global changes in volume appear to be driven by these occipital and parietal regions, as the majority of other regions were largely stable from A.M. to P.M. (**Table 3-1**). Cortical thickness demonstrated the same pattern (**Table 3-2**, **Table 1-3**).

**Figure 3.**
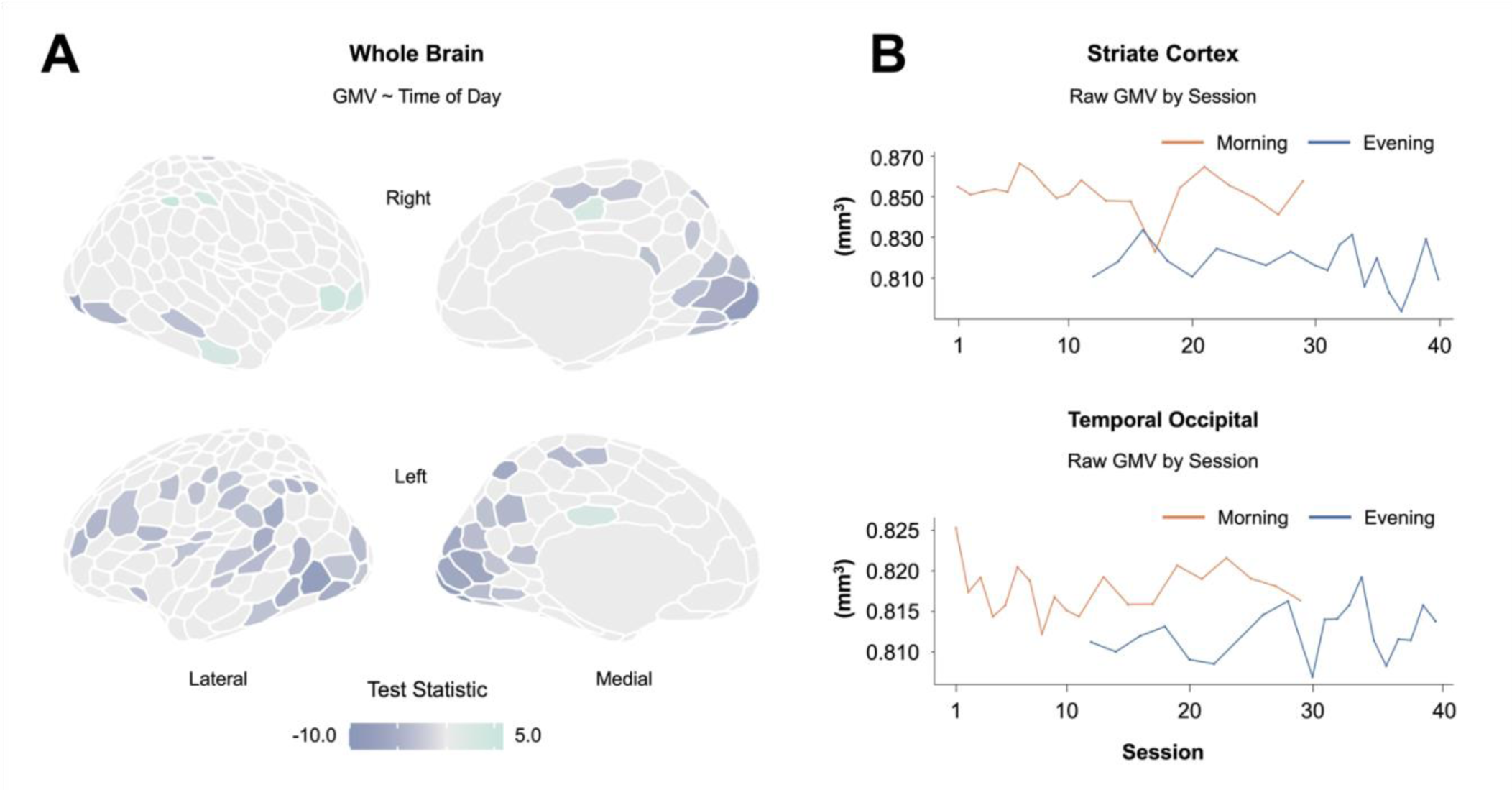
Regional neuroanatomical changes between morning and evening sessions. (**A**) Multivariate regression demonstrates a mostly negative relationship between time of day and gray matter volume (FDR-corrected). **(B)** Gray matter volume in the striate cortex and temporal occipital regions are heightened in the morning and reduced in the evening. Abbreviations: GMV = Gray Matter Volume.

**Table 1.**
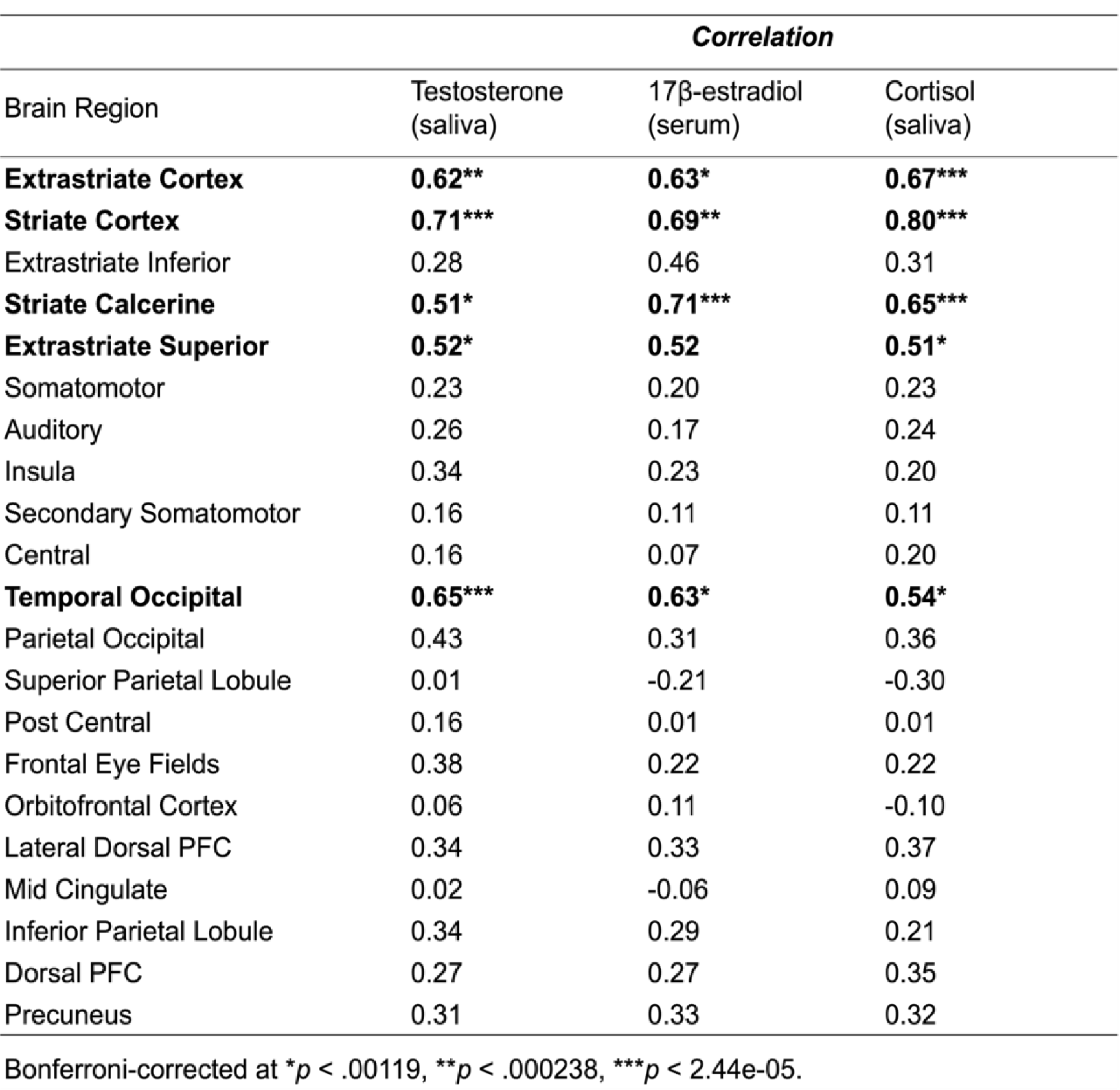
Correlations between GMV in select cortical regions and steroid hormones. Several brain regions, including the extrastriate cortex, striate cortex, striate calcarine, and temporal occipital, were positively correlated with most steroid hormones (testosterone, estradiol, and cortisol). For the full list of cortical correlation results, see **Table 1-2**.

### Diurnal subcortical volume reduction is regionally specific

Total right hippocampal volume decreased significantly from morning to evening (*t*(36.89) = – 5.17, *p* < .001). Similarly, right and left cerebellum and brain stem volumes were significantly reduced from morning to evening scans (*t*(35.59) = –6.41, *p* < .001, *t*(36.27) = –8.38, *p* <.001, and *t*(34.06) = –3.87, *p* < .001, respectively). The effect size was large across these regions (all *d* < –1.24) and they demonstrated positive correlations with steroid hormones (**Figure 4A**, **Table 4-1**). When assessing subfield volumes of the hippocampal body, we observed no changes by TOD. All seven subfield volumes remained stable throughout the day (Bonferroni corrected *p* > .007), and there were no significant relationships between subfield volumes and steroid hormones (**Figure 4B**, **Table 4-2**).

**Figure 4.**
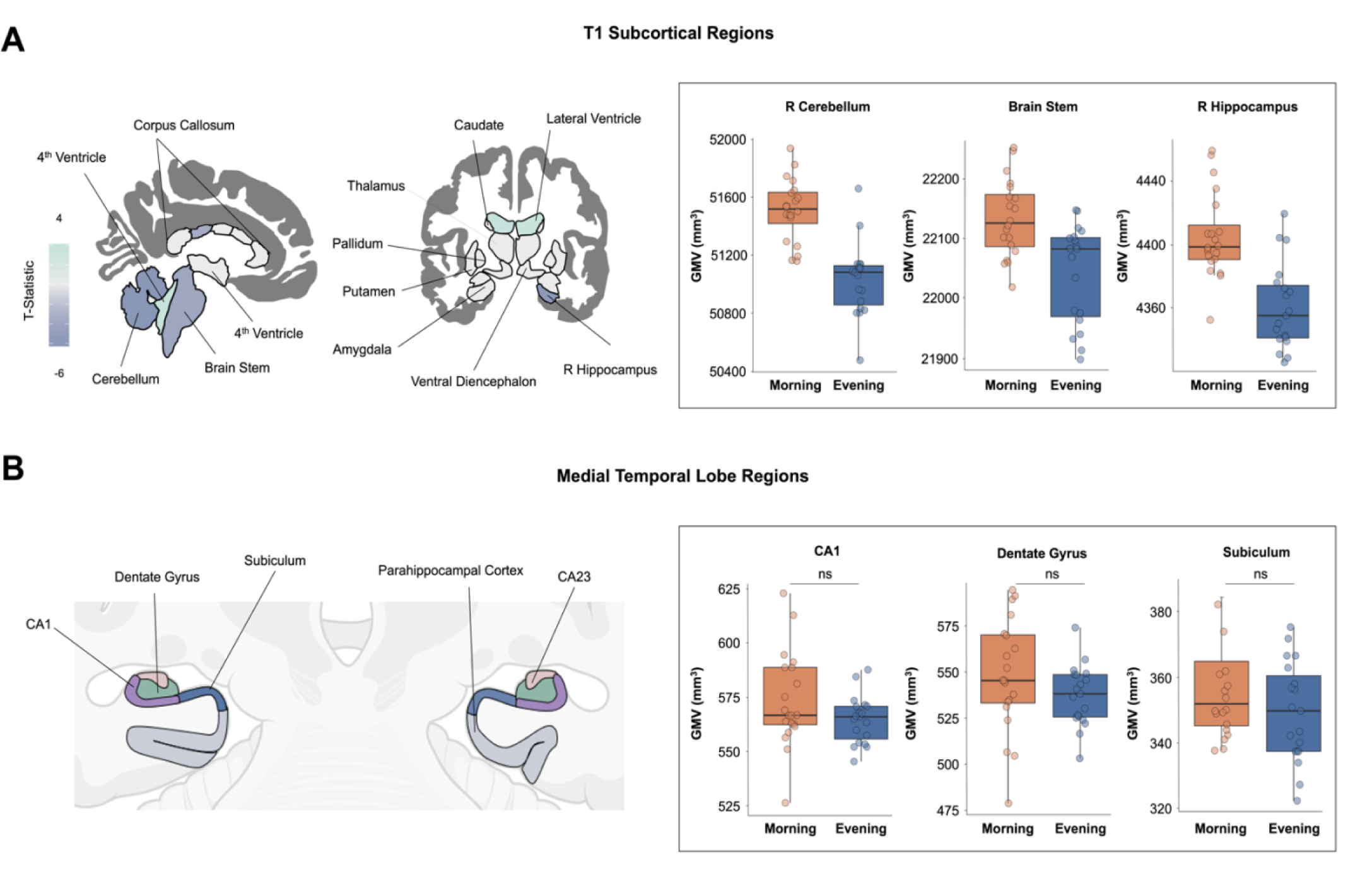
Subcortical gray matter volume by time of day. (**A**) Multivariate regression for subcortical regions demonstrated decreasing volume throughout the day. Left: blue tones indicate volumetric reduction while aqua tones indicate volumetric expansion from A.M. to P.M. (FDR at *q* < 0.05; non-significant test statistics were set to zero for interpretability). Gray matter volume was lower in the right cerebellum, brain stem, and right hippocampus in morning compared to evening sessions. For the full list of subcortical results, see **Table 4-1**. **(B)** Cartoon illustrates medial temporal lobe subregion segmentation from the ASHS parcellation. Gray matter volume in CA1, dentate gyrus, and subiculum did not change throughout the day. This pattern held for all medial temporal lobe subregions (CA2/3, entorhinal cortex, perirhinal cortex, and parahippocampal cortex, *p* > .007 for all, **Table 4-2**). Abbreviations: GMV = Gray Matter Volume, ns = not significant.

## Discussion

Here we investigated the impact of circadian rhythms on global and regional brain morphology in a densely-sampled male. Total brain volume, GMV, and CT decreased from morning to evening, while CSF and ventricle size increased. These changes were not uniform across the cortical mantle, but driven by A.M. to P.M. decreases in GMV and CT in occipital and parietal cortices. These results build on a growing literature demonstrating the robust influence of biological rhythms on human brain morphology.

The direction and magnitude of the observed changes in global brain morphology are consistent with existing MRI studies of TOD. Previous reports suggest a ∼2% decrease in intracranial volume (Trefler et al., 2016) and ∼1% decrease in GMV (Karch et al., 2019) over the course of a day in healthy populations. These changes are on par with the average 0.62% decrease in GMV observed from morning to evening in the present study.

The most prominent changes in brain morphology were evident in parietal and occipital cortices, which showed decreases in both GMV and CT from morning to night. These changes were strongly correlated with steroid hormone levels, including testosterone, estradiol, and cortisol. These regional changes are in keeping with a recent study exploring the effect of diurnal rhythms on brain structure, where GMV and CT in primary visual cortex were tied to diurnal variation in mood and cognition (Zareba et al., 2023). These changes in volume in posterior cortices are mirrored by changes in functional coherence. In a parallel analysis, our group examined changes in functional coherence from A.M. to P.M. in the same densely sampled male. Functional coherence in posterior visual networks were particularly sensitive to TOD. Nodes belonging to central and peripheral visual networks demonstrated some of the strongest associations with diurnal variation in testosterone, estradiol, and cortisol (Grotzinger et al., 2023), paralleling the changes in GMV and CT we see here. Hormonal regulation of visual cortex has been demonstrated in rodent studies. For example, ovarian hormone exposure led to decreased cortical neuron number in the primary visual cortex of female rodents (Nuñez et al., 2002), while androgen exposure prevented cell death in visual cortex (Nuñez et al., 2000). One human study demonstrated TOD effects on GMV outside of the parietal and occipital cortices. The authors scanned individuals once between 10 A.M. and 12 P.M. and again between 2 P.M. and 4 P.M. and reported changes in the frontal and temporal lobe (Trefler et al., 2016). This design was outside of the window aligned with peak and trough of diurnal steroid hormone production, a difference from the present study that could be driving these regional discrepancies in cortical GMV findings.

We also determined the extent to which subcortical structures change by TOD, noting significant decreases in GMV in the cerebellum, brain stem, and right hippocampus. Next, we employed high resolution imaging and segmentation of the hippocampal body to assess medial temporal lobe subregions (including dentate gyrus, CA1, CA2/3, subiculum, dentate gyrus, parahippocampal cortex, entorhinal cortex, and perirhinal cortex), revealing stability over time. The discrepancy between whole hippocampal volume and subfield-level volumes within the hippocampal body suggest that the head and tail of the hippocampus may be the loci of diurnal change, though future work needs to be done to clarify these differences. Existing studies on TOD-related subcortical changes are mixed. Shorter photoperiods have been associated with smaller total hippocampal volumes (Miller et al., 2015) and smaller brainstem volumes (Majrashi et al., 2020). In contrast, (Zareba et al., 2024) reported an increase in GMV in the amygdala and hippocampus from morning to evening. The present study, which tracks brain and hormone dynamics at a high temporal resolution, suggests an overall decline in cortical and subcortical GMV from morning to evening—a pattern that mirrors the diurnal decrease in steroid hormones.

While it is clear that brain morphology exhibits circadian change, the role of steroid hormones in shaping diurnal changes in brain volume is not definitive, as results from the present study are correlational. However, biological rhythms over longer timescales establish steroid hormones’ role in shaping brain morphology. Across the ∼28 day menstrual cycle, endogenous fluctuations in progesterone are associated with changes in GMV in the CA2/3, parahippocampal, entorhinal, and perirhinal cortex regions (Taylor et al., 2020); disrupting this hormonal rhythm eliminates these GMV changes. Further, changes in progesterone are tied to cerebellar changes in functional brain network organization (Fitzgerald et al., 2020). Steroid hormone expression in mammalian cerebellum (Pang et al., 2013) and hippocampus (Brinton et al., 2008) is well-established, underscoring the hormonal regulation of these regions. The present findings are consistent with a model in which steroid hormones contribute to morphological changes in these regions at shorter, circadian, timescales.

Brain volume reduction throughout the day may be related to diurnal regulation of metabolic and synaptic machinery via the glymphatic system. Through the glymphatic pathway, metabolic buildup during the day is removed by the exchange of CSF with interstitial fluid. CSF enters the brain, mixes with interstitial fluid and then exits the brain via venous drainage (Hauglund et al., 2020). This process is mediated by aquaporins, channels that are a key pathway for water movement. Aquaporin subtype 4 (AQP4) is a water channel found in astrocytes that helps maintain water homeostasis in the brain (Badaut et al., 2002). Aquaporin activity may be under the control of circadian rhythms and steroid hormones. In mice, localization of AQP4 to the vasculature, and glymphatic influx, peaked during the daytime, and knocking out AQP4 abolished these day–night changes in glymphatic influx (Hablitz et al., 2020). Aquaporins have also been identified in the paraventricular nucleus of the hypothalamus (Jung et al., 1994), a structure that plays an essential role in driving circadian rhythms and steroid hormone production. Circadian patterns of glymphatic activity may be driven by steroid hormones, which modulate aquaporin activity. Testosterone administration increases AQP4 protein levels in astrocytes (Gu et al., 2003). In contrast, estradiol or progesterone treatment decreases AQP4 expression and brain water content (Soltani et al., 2016). A potential mechanism for steroid hormone–mediated brain changes could be that declining steroid hormones throughout the day coincides with increased AQP4 expression and increased CSF and ventricle volume. Since CSF and ventricle expansion or swelling may lead to reductions in brain volume, an aquaporin-mediated increase in fluid dynamics could explain the observed decline in brain volume throughout the day.

Why might these diurnal rhythms in the brain exist? For most species, behavior is heavily organized around a day/night cycle. Diurnal rhythms are ubiquitous in nature and have broad implications for animal cognition and behavior. The consequences of disrupting circadian rhythms is increasingly well-characterized in animal and human studies (Walker et al., 2020). Major depressive disorder, seasonal affective disorder, and bipolar disorder have all been tied to disruptions in circadian cycles (Dollish et al., 2024). Metabolic side effects, such as abnormal glucose metabolism and lower insulin sensitivity, are also well-established consequences of circadian rhythm disruption (Potter et al., 2016), as are memory impairments (Bellfy et al., 2023; Smies et al., 2022). In short, biological rhythms in the brain likely regulate mood and cognition. In future precision imaging studies, tying cyclic changes in regional and global brain volume to changes in behavior warrants investigation.

This precision imaging study mapped neuroanatomical changes across the circadian cycle in a single male with unprecedented temporal resolution. One limitation inherent to precision imaging studies is the inability to generalize findings to a broader population. However, the diurnal rhythms we observed are consistent with findings reported in cross-sectional studies. Future precision imaging studies can expand on this work by sampling demographically diverse individuals, including those with disrupted circadian rhythms (e.g., individuals who work a night shift, flight attendants) and altered hormonal milieus (e.g., via pharmacological manipulation). Such studies would aid in disentangling the unique role of steroid hormones from other variables that may contribute to the rapid structural changes occurring in the brain diurnally. These findings have substantial implications for the neuroimaging community: given the pronounced effects of diurnal rhythms on brain morphology, MRI experiments should adjust for TOD in all analyses.

Finally, hormonal fluctuations in females are often considered a source of unwanted variability– dogma that was invoked to exclude female animals from half a century of biomedical research (Beery & Zucker, 2011; Shansky, 2019).This study aids in efforts to debunk that myth; steroid hormones influence aspects of brain morphology and function in males and females. In all cases, the rhythmic production of steroid hormones should not be considered noise, but rather a foundational feature of biology.

## Author Contributions

Conceptualization: L.P, C.M.T., E.G.J.; Formal analysis: E.M.M.; Funding Acquisition: E.G.J.; Investigation: L.P., Methodology: E.M.M., L.P., C.M.T., E.G.J.; Project Administration: E.M.M., L.P., H.G.; Resources: E.G.J.; Supervision: E.G.J.; Visualization: E.M.M., H.G.; Writing – Original Draft: E.M.M.; Writing – Review and Editing: L.P, H.G., C.M.T., E.G.J.

## Data Availability Statement

The dataset consists of 40 MRI scans (T1w, T2w, and resting-state functional scans) alongside state-dependent measures and serum assessments of steroid hormones for each session. This data will publicly available on https://openneuro.org/ upon publication.

## Acknowledgements

This study was supported by the Ann S. Bowers Women’s Brain Health Initiative (EGJ, CT), UC Academic Senate (EGJ), and NIH AG063843 (EGJ). We thank Mario Mendoza for phlebotomy and MRI assistance.

